# A mechanistic understanding of the effect of *Staphylococcus aureus* VraS histidine kinase single point mutation on antibiotic resistance

**DOI:** 10.1101/2025.01.06.631495

**Authors:** Liaqat Ali, Salima Karki, Gunavanthi D Boorgula, Amir Mekakda, Brittnee Cagle-White, Shrijan Bhattarai, Robert Beaudoin, Aryanna Blakeney, Sanjay Singh, Shashikant Srivastava, May H. Abdelaziz

## Abstract

Bacterial genomic mutations in *Staphylococcus aureus (S. aureus)* have been detected in isolated resistant clinical strains, yet their mechanistic effect on the development of antimicrobial resistance remains unclear. The resistance-associated regulatory systems acquire adaptive mutations under stress conditions that may lead to a gain of function effect and contribute to the resistance phenotype. Here, we investigate the effect of a single-point mutation (T331I) in VraS histidine kinase, part of the VraSR two-component system in *S. aureus.* VraSR senses and responds to environmental stress signals by upregulating gene expression for cell wall synthesis. A combination of enzyme kinetics, microbiological, and transcriptomic analysis revealed the mechanistic effect of the mutation on VraS and *S. aureus*. Michaelis Menten’s kinetics show that the VraS mutation caused an increase in the autophosphorylation rate of VraS and enhanced its catalytic efficiency. The introduction of the mutation through recombineering coupled with CRISPR-Cas9 counterselection to the Newman strain wild-type (WT) genome doubled the minimum inhibitory concentration of three cell wall-targeting antibiotics. The mutation caused an enhanced *S. aureus* growth rate at sub-lethal doses of the antibiotics, confirming the causative effect of mutation on bacterial persistence. Transcriptomic analysis showed a genome-wide alteration in gene expression levels and protein-protein interaction network of the mutant compared to the WT strain after exposure to vancomycin. The results suggest that *vraS* mutation causes several mechanistic changes at the protein and cellular levels that favor bacterial survival under antibiotic stress and cause the mutation-harboring strains to become the dominant population during infection.

**Importance:** Rising antimicrobial resistance (AMR) is a global health problem. Mutations in the two-component system have been linked to drug- resistance in *Staphylococcus aureus*, yet the exact mechanism through which these mutations work is understudied. We investigated the T331I mutation in the *vraS* gene linked to sensing and responding to cell wall stress. The mutation caused changes at the protein level by increasing the catalytic efficiency of VraS kinase activity. Introducing the mutation to the genome of an *S. aureus* strain resulted in changes in the phenotypic antibiotic susceptibility, growth kinetics, and genome-wide transcriptomic alterations. By a combination of enzyme kinetics, microbiological, and transcriptomic approaches, we highlight how small genetic changes can significantly impact bacterial physiology and survival under antibiotic stress. Understanding the mechanistic basis of antibiotic resistance is crucial to guide the development of novel therapeutic agents to combat AMR.

## Introduction

*Staphylococcus aureus* (*S. aureus*) is a major pathogenic bacterium that causes serious skin, soft tissue, and nosocomial infections that could progress to severe illnesses, such as osteomyelitis, endocarditis, sepsis, and bacteremia in humans (1). Methicillin-resistant *S*. *aureus* (MRSA) is the leading cause of nosocomial infections in hospitals and community settings (2). In 2020, during the COVID-19 pandemic, the rate of MRSA infections in the United States increased by 13% compared to that reported during 2017 – 2020 (3). The Infectious Diseases Society of America MRSA treatment guidelines recommend either vancomycin or daptomycin as first-line agents (4).

The MRSA phenotype is known to arise as a result of acquired external genetic elements of resistance genes like *mecA* (5). The vancomycin-intermediate *S. aureus* (VISA) is characterized by vancomycin minimum inhibitory concentration (MIC) ranging between 4 – 8 µg/mL. Whole genome sequencing revealed that the VISA phenotype may arise due to mutations within various loci of the *S. aureus* genome. These mutations are mainly adaptive responses driven by environmental stress, including antibiotic exposure (6–8). In addition, mutations in VISA have also been reported in the two-component systems (TCSs) network (9, 10).

The network of TCSs has several sensory roles in bacteria, including cell wall biosynthesis, such as VraSR, WalKR, and GraSR (11–13). These systems initiate a response mostly after exposure to antibiotics that target the cell wall. For instance, *S*. *aureus* utilizes VraSR (vancomycin resistance associated two-component system) to respond to glycopeptides or beta-lactam antibiotics activity (14). The VraS histidine kinase is a transmembrane receptor that phosphorylates the response regulator VraR which then binds to bacterial DNA and activates *vraSR* regulon and several other genes, including *pbp2*, *murZ*, *sgtB*, *fmtA*, and *lytR*, whose products collectively overcome the activity of cell wall inhibiting antibiotics (15, 16). Previous studies show that vancomycin exposure can upregulate VraSR expression, resulting in enhanced intracellular survival of *S. aureus* (15, 17, 18). A number of point mutations in *vraSR* regulon have been reported to be associated with VISA (10, 19–22).

Structurally, VraS belongs to the intramembrane-sensing histidine kinases family that consists of a transmembrane domain, dimerization interface with a conserved histidine residue, and an ATP-binding catalytic domain (23). Among these domains, the ATP-binding domain is implicated in the autophosphorylation of VraS (24). Mutations in this domain were linked to increasing resistance to antibiotics (10, 21). A recent study by Sabat et al revealed that prolonged daptomycin treatment of infective endocarditis with susceptible *S*. *aureus* led to the emergence of a daptomycin-resistance and whole genome sequencing revealed a VraS T331I mutation in the clinical isolate (22). To further expand the understanding of VraS mutations and the associated mechanism of antibiotic resistance, we performed a series of experiments targeting this T331I mutation located in the ATP-binding domain of VraS using a combination of biochemical, microbiological, and transcriptomic techniques. By unraveling the consequences of the mutation on VraS catalytic functions and *S. aureus* at the cellular level, we attempt to uncover the mechanistic basis for developing antibiotic resistance in that highly pathogenic organism.

## Results

### VraS T331I mutation causes an increased autophosphorylation rate and catalytic efficiency

The VraS wild-type (WT) construct was amplified from the previously described pGEX- 4T-1 plasmid (25). The gene was inserted into a pET15b plasmid and modified to introduce a single point mutation at the T331 residue as described in the Material and Methods section. The constructs were expressed in BL21 (DE3) pLysS strain and purified using Ni-NTA affinity and gel filtration chromatography (Figure 1 and S1). The identity of the proteins was confirmed by Western blotting with anti-His antibody, and a coupled kinase kinetic assay was used to assess the autophosphorylation reaction rate of VraS WT and T331I, as previously described (26). The reaction rate for T331I was 223.73 ± 5.6 nmol hr^-1^ compared to 19.03 ± 1.1 for the WT enzyme, a ∼12-fold increase (Figure 2A). Using increasing ATP concentrations, we performed Michaelis Menten’s kinetic analysis. The calculated ATP affinity to WT VraS (*K_M_*) was 5.34 ± 1.9 µM compared to 7.95 ± 0.1 µM for the T331I mutant, which was not a significant change. However, the catalytic rate of the reaction (*k_cat_*) increased from 32.02 ± 3.7 to 170.70 ± 2.3 hr^-1^, causing the overall catalytic efficiency (*K_M_* /*k_cat_*) of the mutant to increase more than 3-folds from 6.60 ± 1.7 to 21.37 ± 0.3 µM^-1^ hr^-1^ compared to WT VraS. The results can be attributed to the location of the T331 residue in the catalytic binding site in close proximity to the bound ATP (Figure 2B).

**Figure 1:**
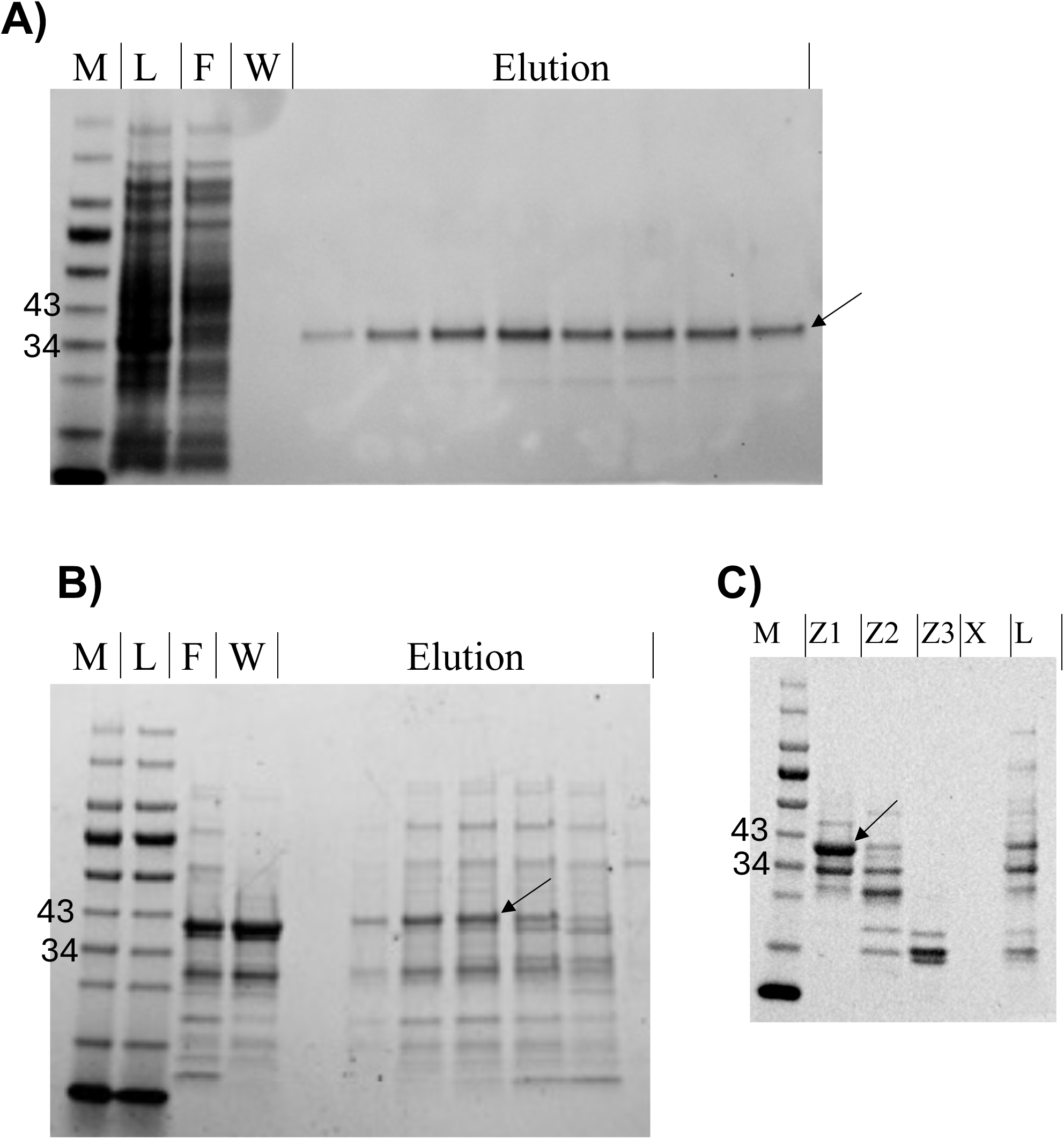
Purification of VraS WT and T331I mutant. SDS-PAGE images of A) VraS WT fractions and B) T331I mutant fractions from the His-Trap column, and C) T331I mutant fractions from the gel filtration column (M: molecular weight standard ladder, L: load pre-column, F: flowthrough, W: wash, Elution represent fraction in the protein eluted peak, Z1-3: representative fractions from the gel filtration column).

**Figure 2.**
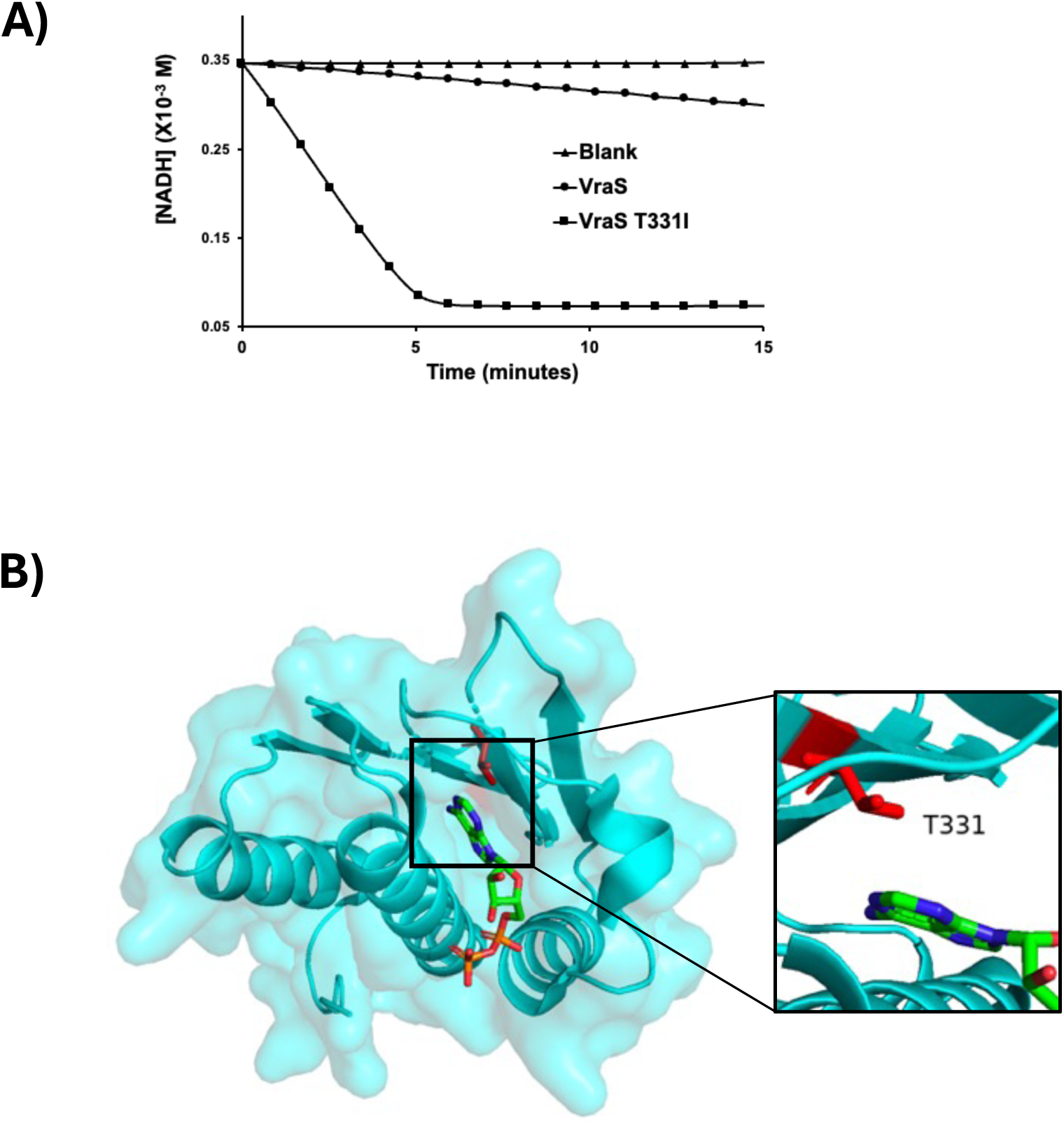
Autophosphorylation of VraS. A) The autophosphorylation reaction rate of VraS WT and T331I mutant (5 μM) as a function of the rate of NADH disappearance, compared to a blank reaction. The reaction reaches a plateau with the T331I as the NADH in the assay mixture is depleted. B) Surface representation of the crystalized VraS catalytic domain (PDB 4GT8). The inset shows T331 (red sticks) proximity to the bound ATP (in colored sticks).

### Shift in minimum inhibitory concentration (MIC) and drug’s efficacy and potency in the VraS T331I mutant

The T331I mutation was introduced to the genomic background of the *S. aureus* susceptible Newman WT strain using recombineering techniques with CRISPR-Cas9 counterselection per published protocols (27). Broth microdilution MICs were performed with vancomycin, methicillin, and daptomycin (in the presence of 50 mg/L CaCl_2_) for the WT and the T331I mutant strains in two growth media, cation-adjusted Muller Hinton broth (CAMHB) and Lysogeny Broth (LB). The MICs for the T331 mutant show a 2-fold increase with most of the tested antibiotics (Table 1). Next, concentration-response studies with the three antibiotics were performed using both WT and mutant strains. Data was analyzed using the inhibitory sigmoid maximal effect model to determine the relationship between the drug concentrations and the bacterial burden and is reported here as mean ± standard error **(Figure 3 and Table 1 and S1)**. Two-way analysis of variance did not show significant differences in the vancomycin efficacy (E_max_ or maximal effect) and potency (EC_50_ or drug concentration mediating 50% of the E_max_). However, the EC_50_ of the mutant strain was higher for methicillin and daptomycin compared to the WT strain, indicating the possible role of the *vraS* mutation in antibiotic resistance.

**Figure 3.**
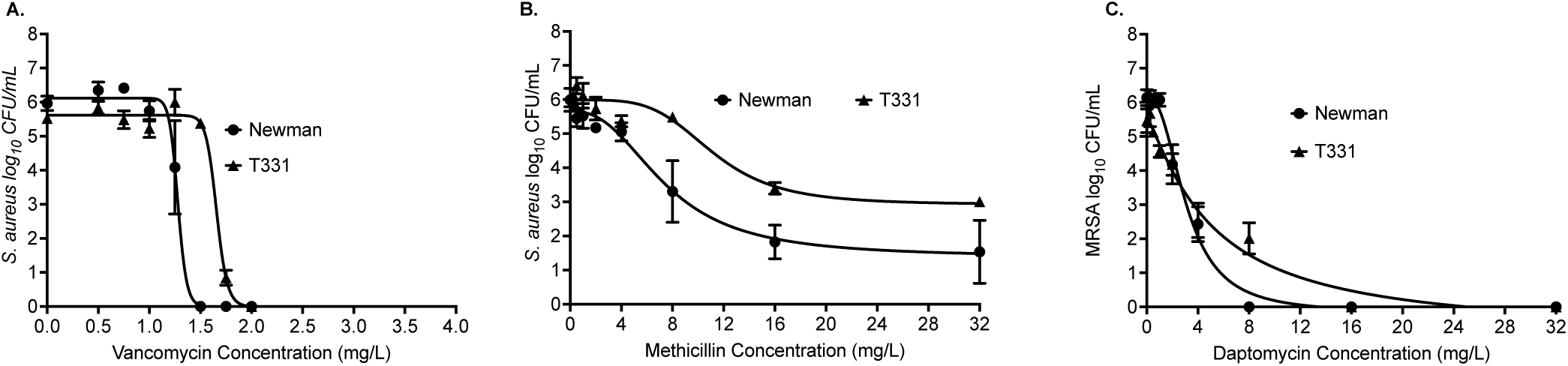
Relationship between the drug concentrations and the bacterial burden in the Newman WT and T331I mutant strain.

**Table 1.**
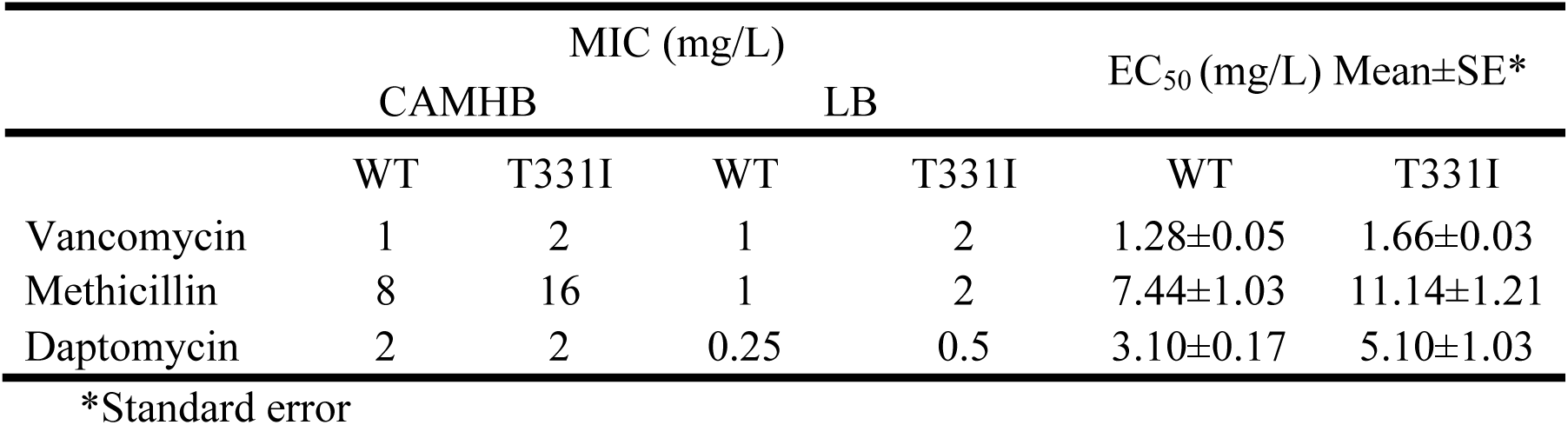
The effect of the mutation on drug susceptibility using different media.

| | MIC (mg/L) | | | | $EC_{50}$ (mg/L) Mean±SE* | |
| --- | --- | --- | --- | --- | --- | --- |
|  | CAMHB |  | LB |  | WT | T331I |
|  | WT | T331I | WT | T331I |  |  |
| Vancomycin | 1 | 2 | 1 | 2 | 1.28±0.05 | 1.66±0.03 |
| Methicillin | 8 | 16 | 1 | 2 | 7.44±1.03 | 11.14±1.21 |
| Daptomycin | 2 | 2 | 0.25 | 0.5 | 3.10±0.17 | 5.10±1.03 |

\*Standard error

### The T331I strain demonstrates significant changes in *S. aureus* growth kinetics under antibiotic stress

In the presence of resistant mutant strains of pathogens, the window of concentration between MIC of the WT to that required to inhibit the least susceptible mutant is expected to enrich the resistant strain subpopulations selectively (28). To assess the extent of this window with the T331I mutation, the growth curves of WT and mutant strains were monitored with and without antibiotic stress, as previously described (25). The strains were grown for 24 hours in the presence of different doses of vancomycin, daptomycin, and methicillin. The antibiotic concentrations screened were below to slightly above the MIC for the strains. In the absence of antibiotics, the T331I strain had similar growth kinetics to the WT strain with no significant growth defect. The calculated growth rate for WT Newman was 0.09 (95% CI: 0.08 to 0.10) log_10_ CFU/mL/h, whereas the T331I was 0.07 (95% CI: 0.06 to 0.8) log_10_ CFU/mL/h. However, in the presence of antibiotics, the mutant shows faster growth rates, shorter lag time, and more persistence than the WT strain (Figure 4).

**Figure 4.**
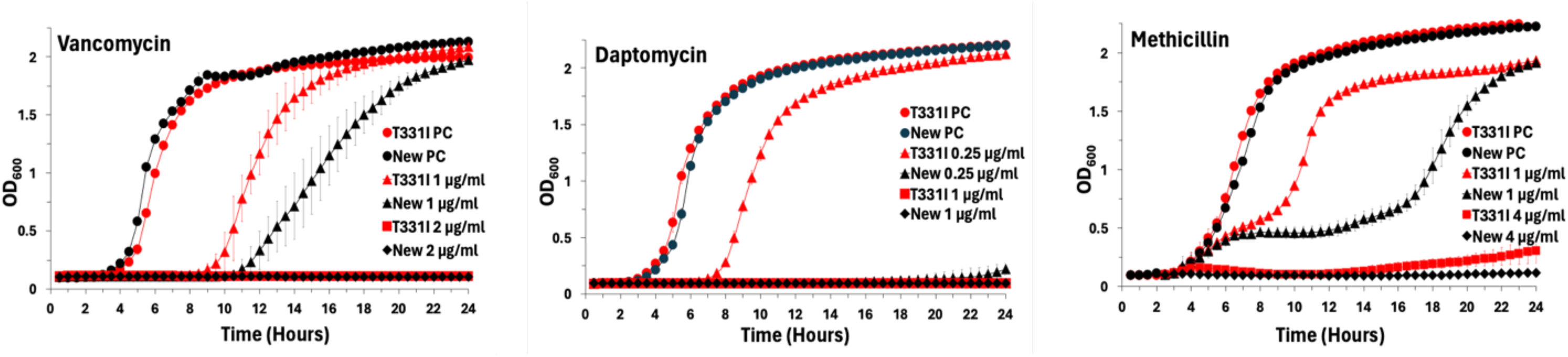
Growth curves of the T331I mutant (red lines) and WT (New) strains (black lines) in the presence of different antibiotic concentrations. PC indicates positive control growth without antibiotics. The data represent the average ± SD of at least three biological replicas.

### The expression of *blaZ* is significantly increased in the T331I mutant strain under antibiotic stress

To investigate the possible effect of mutation on VraSR self-regulation and its downstream transcription, we compared the expression of *vraR*, *pbp2*, and *blaZ* in the WT and T331I strains before and after antibiotic stress using quantitative real-time PCR analysis (qRT-PCR), as previously described (25) and detailed in the Materials and Methods section. The *pbp2* gene translates to a transpeptidase that is essential for cell wall formation and highly involved in resistance to antibiotics, while *blaZ* translates to the β-lactamase enzyme responsible for the hydrolysis of β-lactam antibiotics (29). Figure 5 shows that in the absence of antibiotic stress, T331I had similar expression levels to the WT strain for *vraR* and *pbp2* yet showed a 6.5-fold increase in *blaZ* expression (*p*= 0.0209). After vancomycin exposure, the two strains showed a similar trend of increased expression for the tested genes. Compared to the WT, T331I showed a 1.7-fold lower *pbp2* expression (*p*<0.0001) but a 2.5-fold higher *blaZ* expression (*p*=0.0001). Methicillin caused an increase only in *blaZ* expression with T331I showing 3.5-fold higher expression than WT (*p*<0.0001). Interestingly, daptomycin did not affect the expression level of the tested genes in the WT with all antibiotics. In comparison, T331I had significantly higher expression of *pbp2* (1.5-fold, *p*=0.0224) and *blaZ* (6.5-fold, *p*=0.0027) compared to the WT strain.

**Figure 5.**
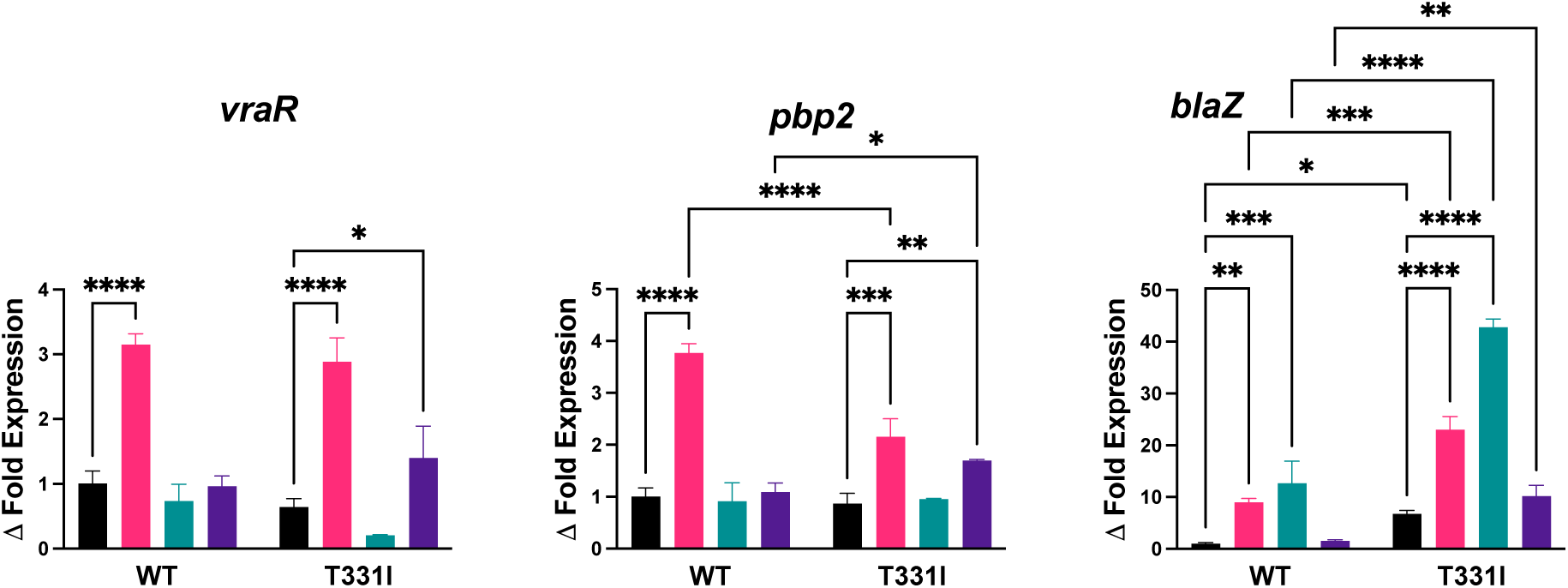
Fold change in expression levels of *vraR*, *pbp2,* and *blaZ* after mutation and/or antibiotic exposure (black: no antibiotic control, magenta: vancomycin, green: methicillin, and purple: daptomycin). The data represent the mean ± SE (*n* = 3), and statistical significance between control versus treated samples for each strain and between WT and T331I under the same treatment was calculated via one-way ANOVA with multiple comparisons.

### Significant transcriptomic changes of the T331I mutant in response to vancomycin exposure

We performed RNA-seq to determine the transcriptomic changes in the T331I mutant compared to the Newman WT before and after vancomycin exposure (2X MIC for 1 hour). In the absence of stress, T331I had 70 differentially expressed genes (DEGs), with 22 genes significantly dysregulated by >2-fold compared to the WT strain, representing 0.7% of the detected *S. aureus* genome (Table 2, Figure 6A). The WT strain showed 287 DEGs in response to vancomycin exposure, including 37 dysregulated genes by >2-fold change (Table 2, Figure 6B). Subjecting the T331I mutant strain to the same antibiotic stress resulted in large-scale changes in transcription with >1200 DEGs compared to T331I in the absence of stress or WT in the presence of the same stress, where in each case >450 genes (∼14% of the genome) were dysregulated by >2-fold change and ∼3% by >4-fold change (Table 2, Figure 6C and 6D). This represents more than a 10-fold change in genome-wide gene expression in response to vancomycin for the mutant strain compared to WT (the lists of genes can be found in supplementary material). The gene expression profile shows that several genes were uniquely expressed in each strain, with and without vancomycin stress (Figure S2 and supplementary material).

**Figure 6.**
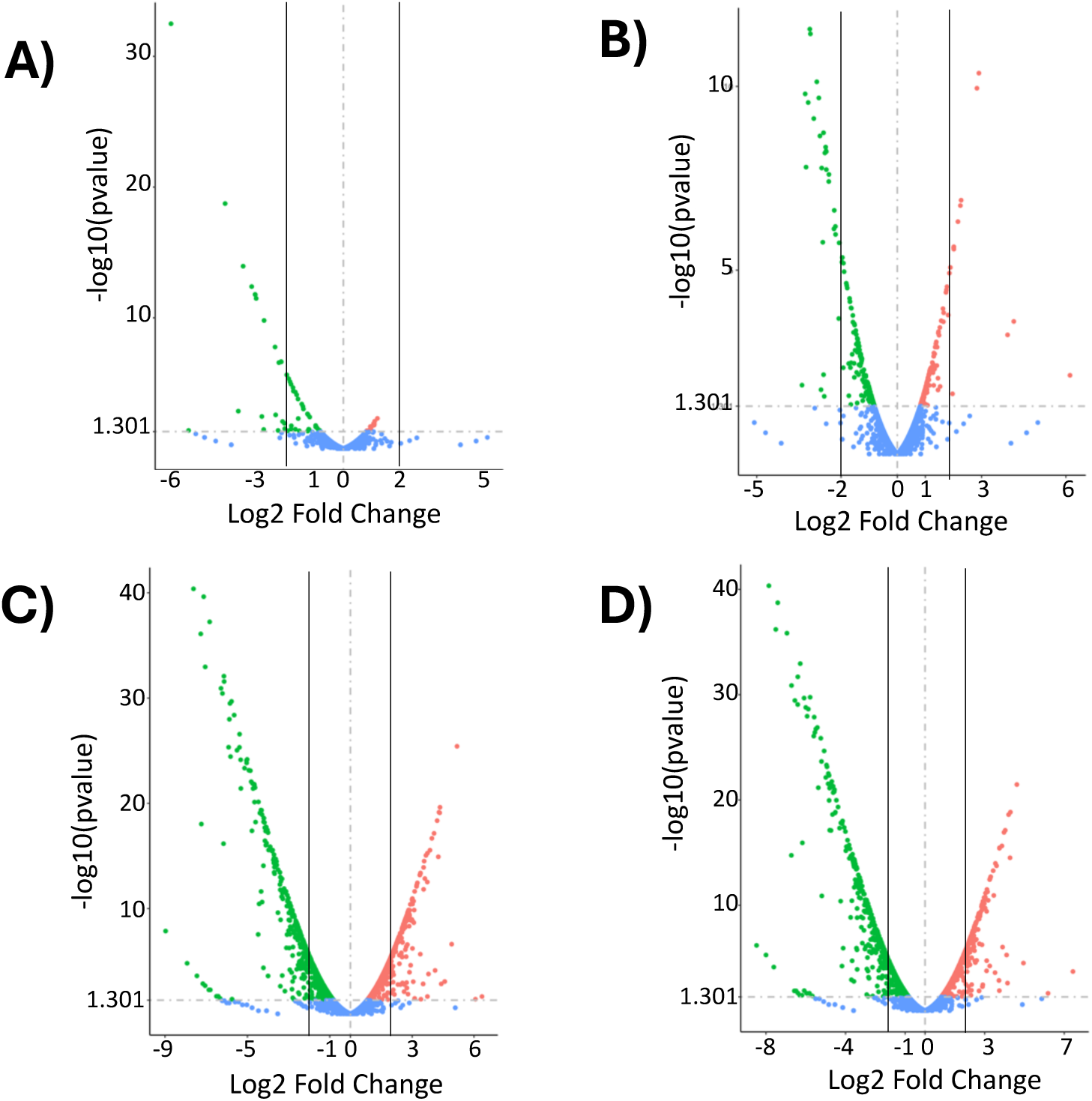
Volcano plots of differential gene expression between T331I and WT strains with and without vancomycin stress. A) T331I compared to WT, B) WT with vancomycin compared to WT, C) T331I with vancomycin compared to T331I, and D) T331I with vancomycin compared to WT with vancomycin. Colored dots (red = upregulated, green = downregulated, and blue = not significantly changed) based on *p*-value (< 0.05) and log2(fold-change) and logFC cutoff of 1.3. The vertical lines represent the 2-fold change in gene expression selected to generate the data in Table 2.

**Table 2.**
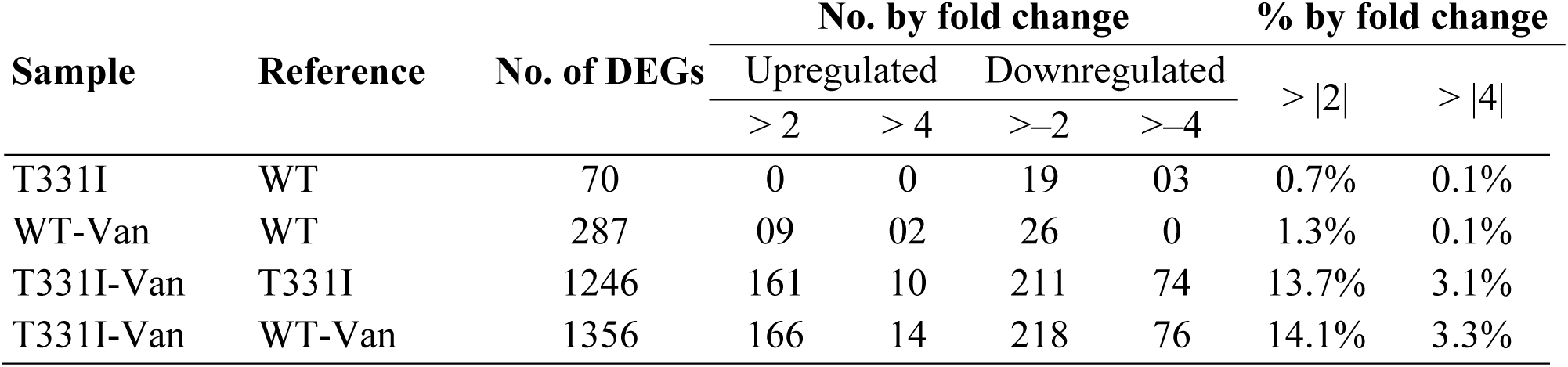
Comparison of differentially expressed genes (DEGs) in WT and T33I mutant strains (genes are listed in supplementary material).

To verify the RNA-seq results, qRT-PCR was performed on the top 4 dysregulated genes to confirm the change in expression quantitively using primers listed in Table S2. The RNA-seq results show that *treP* and *isaA* genes were 5.2 and 4.4-fold upregulated; respectively, in the T331I mutant versus WT under vancomycin stress, while the *hisD* and *cwrA* genes were 7.3 and 6.8-fold downregulated. The dysregulated gene expression trends observed in qRT-PCR were similar to those captured in RNA-seq (Figure S3). To determine the biological functions of the DEGs in T331I versus WT after vancomycin exposure, a Kyoto Encyclopedia of Genes and Genomes (KEGG) pathway enrichment analysis was performed (30). Pathways involved in the ribosome and amino acid degradation were significantly upregulated. The down-regulated pathways include the biosynthesis of amino acids, biosynthesis of secondary metabolites, and 2- oxocarboxylic acid metabolism. The pathway of microbial metabolism in diverse environments was highly dysregulated, with 41 upregulated and 48 downregulated genes (Figure S4).

### The T331I mutation causes significant changes in *S. aureus* predicted protein–protein interaction (PPI) network

The STRING analysis can predict the PPIs that occur in a strain compared to its control which translates the alterations in gene expression to the proteomics level. The STRING database (http://string-db.org/) was used for the analysis of predicted PPIs to understand the effect of T33I mutation with and without vancomycin exposure and visualized using Cytoscape 3.10.1 software (31). As a result of the antibiotic stress on the WT strain, a total of 406 PPIs were observed, where VraS showed interactions with VraR, LiaF (VraT), TcaA, and a hypothetical protein (CNH35_RS10705), these constitute VraS “core interactions” (Figure 7A). In the absence of antibiotics, the T331I mutant showed an additional 28 PPIs compared to the WT strain. VraS in the T331I mutant was able to retain its core interactions, while VraR showed two additional interactions with FmtA (teichoic acid D-Ala esterase) and TDCP (transglycosylase domain-containing Protein) (Figure 7B). When the T331I mutant was exposed to vancomycin, the PPIs network became more complex with an additional 4273 PPIs, a 10-fold increase compared to WT under the same antibiotic stress. Along with its core interactions, VraS showed two additional PPIs with GraR and NreC, while VraR showed three additional interactions with GraR, HssS, and NreB, components in other *S. aureus* TCS signaling network (Figure 7C). The results suggest that the mutation may have caused a signal crosstalk between VraSR and other two-component systems only under vancomycin stress to facilitate bacterial survival.

**Figure 7.**
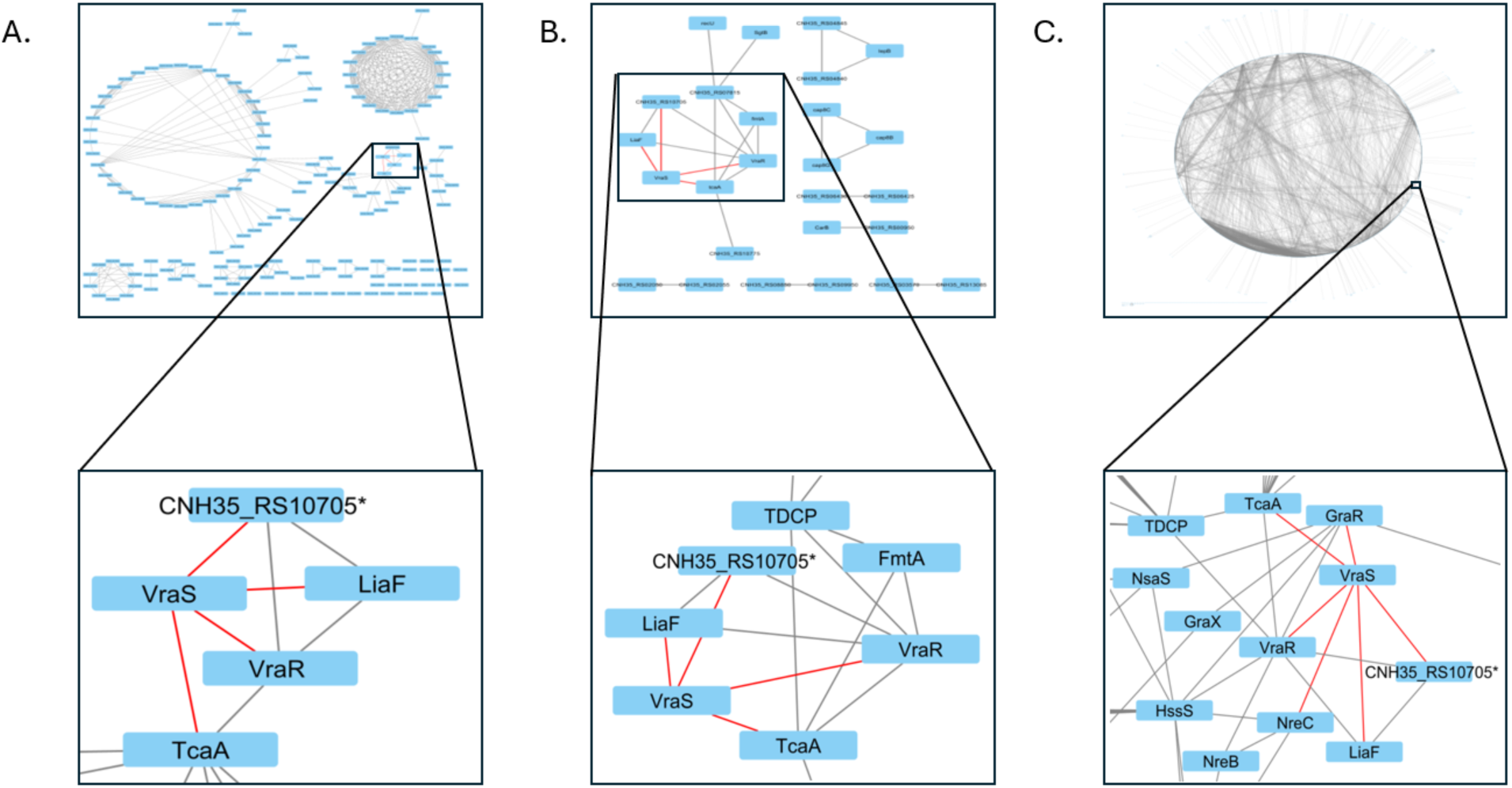
The predicted change protein-protein interaction (PPI) network in *S. aureus* after mutation and/or vancomycin stress as visualized using Cytoscape 3.10.1. The inset shows a Zoomed view of the interactions of the VraSR system, and the red lines highlight VraS PPIs. A) PPIs for WT strain after exposure to vancomycin, B) PPIs for T331I compared to WT, and C) PPIs of T331I after vancomycin compared to T331I.

## Discussion

In *S*. *aureus*, VraSR is activated when the bacteria is exposed to external antimicrobials (15). Previous studies have shown that point mutations in *vraS* tend to show up in strains with more resistance against cell wall-inhibiting antibiotics such as vancomycin, teicoplanin, and oxacillin (10, 21, 32–36)Our results show that the T33I mutation affects VraS catalysis, as a 12- fold increase in autophosphorylation rates compared to the WT enzyme was reported, with a 3- fold increase in the overall catalytic efficiency. The T331I mutation is in close proximity to the ATP binding site located in the VraS catalytic domain. Mutations in the ATP-binding region can induce significant changes in enzymatic activity and ATP consumption, resulting in increased response regulator phosphorylation levels and ultimately upregulating the cell wall synthesis gene cluster and other regulatory genes (37, 38).

The introduction of T33I mutation to the antibiotic susceptible Newman strain genome resulted in increased MICs of the three chemically diverse antibiotics used in the study that affect the cell wall and cell membrane. The growth kinetic curves, shown here, support the presence of a mutant selection window where the mutated strain’s subpopulation (drug-resistant) has the potential to dominate the infection site and enhance bacterial persistence in the presence of drugs. Notably, both the WT and mutant strains exhibited a biphasic growth curve under methicillin treatment. With the WT, the decreased initial growth followed by the surge after 16 hours could be attributed to the lower stability of methicillin in aqueous media (39). Yet the T331I mutant follows the same pattern, with the surge occurring much earlier after 8 hours. The growth profile can thus be explained by the increased expression of the *blaZ* lactamases (supported by the qRT-PCR results) much earlier in the mutant strain that hydrolyzes the antibiotic and restores the bacterial growth rates.

The genes encoding resistance mechanisms vary depending on how they are acquired, either on mobile genetic elements or encoded in the bacterial chromosome. *S. aureus* exhibits two prominent resistance mechanisms: beta-lactamase production and PBP2 production (40). The expression levels of *vraSR* and other resistance-related genes are known to significantly increase after exposure to cell wall-targeting antibiotics (15, 41). Mutations in the VraS lead to an increased expression of *vraSR* regulon and other cell wall synthesis and resistance-related genes (42). In the absence of antibiotic stress, our qRT-PCR results indicated that the WT and T331I mutant strains did not show a difference in expression for the tested genes. The gene expression levels of *pbp2* and *vraR* were significantly increased in both WT and T331I after vancomycin exposure; however, the effect was not observed in WT with daptomycin or methicillin. These findings agree with previously published data with carbenicillin (25). The T331I strain showed a robust response to daptomycin for both *vraR* and *pbp2* increased expression, which may explain the resistance to daptomycin in the clinical isolate harboring the T331I mutation. Concurrently, we found significant changes in *blaZ* expression for T331I upon exposure to all three studied antibiotics compared to the WT strain explaining its resistance to beta-lactam antibiotics.

Under vancomycin stress, the RNA-seq data indicated a wide-scale alteration in gene expression in the T331I mutant compared to the WT strain, which highlights that *S*. *aureus* responded differently in the presence of antibiotic exposure. The differential gene expression was spread over several pathways, including ribosome and peptidoglycan biosynthesis, and was activated in T331I relative to that in WT. Our results, in agreement with previous findings, indicate that the VraS mutation plays an important role in turning on different biological pathways related not strictly to cell wall upregulation but also to bacterial metabolism that significantly impacts antimicrobial susceptibility (20, 43). Therefore, further studies are warranted to determine the role of VraS point mutations in the metabolic pathways we identified to better understand the impact of VraSR on *S. aureus* physiology.

Among the top differentially expressed genes, we found that *treP* and *isaA* were significantly upregulated in T331I, especially after the vancomycin treatment. The TreP facilitates the conversion of trehalose into trehalose-6-phosphate and correlates with transient drug tolerance and inducing resistance (44). IsaA, on the other hand, is involved in peptidoglycan hydrolysis (45) and its deletion led to a significant decrease in β-lactam resistance and biofilm formation (46). The upregulation of *treP* and *isaA* in the T331I mutant suggests their possible involvement in the resistant phenotype of the mutant. The genes associated with the histidine biosynthesis pathway (including *hisD* and *hisB)* as well as *cwrA* were significantly downregulated in T331I. Amino acids such as histidine, arginine, leucine, and valine are important for the survival of *S. aureus* (47), and the decrease in histidine biosynthesis may correlate with altered metabolism in the mutant. On the other hand, *cwrA* is a VraR-regulated gene encoding a cell wall inhibition-responsive protein, and the decrease can attenuate its negative feedback mechanism on cell wall synthesis.

The PPI network shows predicted interactions between VraS, VraR, and VraT (a well-known accessory protein for the VraSR system) (48). The *tcaA* and *fmtA* genes are under the transcription control of VraR, and the results imply an interaction may occur between VraSR and these genes in WT under vancomycin stress and in the T331I mutant regardless of stress. The predicted interaction with TDCP in T331I can result in an increase in the overall transglycosylase activity, which may explain the mutant’s decreased level of *pbp2* expression as one of the major transglycosylase enzymes in *S. aureus* (49). The vancomycin stress on T331I caused additional interactions to appear for VraSR with other two-component systems, GraSR, HssSR, and NreBC, which implies enhanced co-expression and a concerted response to enhance the mutant’s persistence through possible crosstalk that may occur in the signaling network (36).

In conclusion, the T331I mutation in the VraS ATP binding domain enhances resistance against glycopeptides and beta-lactam antibiotics through several underlying mechanisms, such as increased VraS catalytic ability and large-scale alterations in gene expression across multiple bacterial pathways, including resistance-related genes. Targeting the VraS ATP-binding domain could be an effective strategy against drug-resistant *S. aureus* infections. Since mutations in the VraSR system led to antibiotic resistance, there is a need to develop specific inhibitors that target these mutant forms to overcome the problem of drug resistance in *S. aureus*.

## Materials and Methods

### Strains, Media, and Reagents

All chemicals were obtained from Fisher Scientific unless otherwise stated. Tris-Glycine-SDS was from Invitrogen, Mini-PROTEAN TGX stain-free gel and unstained protein standard ladder were from Biorad. *NdeI and XhoI* restriction enzymes, Instant Sticky-end Ligase Master Mix, and Q5 High-Fidelity DNA Polymerase were from NEB. The drug-susceptible strain *S. aureus* Newman D2C strain (ATCC #25904) and its isogenic mutant VraS T331I were maintained in Luria-Bertani (LB) media at 37 °C unless otherwise mentioned. The *Escherichia coli* strain BL21 (DE3) pLysS (Invitrogen #C606010) was used for bacterial heterologous expression of the constructs (all strains used are listed in Table S3).

### Plasmids for Protein Expression

To generate the His-tagged VraS construct, the previously described VraS-KD fragment was amplified from the pGEX-VraS plasmid (25). Primers 1&2 used were designed for amplification to add an *NdeI* restriction site upstream of the VraS-KD gene. The resultant fragment and pET15b bacterial expression plasmid were treated with *NdeI/XhoI* restriction nucleases for 1 hour at 37 °C, then ligated using the Ligase Master Mix. The T331I mutation was introduced by site-directed mutagenesis with Q5 high-fidelity polymerase through the generation of two amplified fragments with overlap at the T331I codon switch (primers 3&4) following the manufacturer’s protocol. The parent template was removed with DpnI (methylation-dependent endonuclease) treatment, and DH5α cells were transformed with the generated plasmids (pET15b-VraS and pET15b-VraS T331I). Plasmids were then isolated from the positive transformants, and the desired modifications were confirmed by sequencing (primers and oligos are listed in Table S2, plasmids are listed in Table S4).

### Large-scale heterologous bacterial expression and purification

The pET15b-VraS and pET15b-VraS T331I plasmids were transformed into BL21 (DE3) pLysS cells and incubated overnight at 37 °C on LB plates supplemented with ampicillin (100 µg/ml). Positive transformants were cultured in 50 mL of terrific broth (TB) supplemented with ampicillin (100 µg/ml) overnight at 37°C, then subcultured to 1 Liter the next day with shaking at 200 rpm. The cultures were induced at mid-log growth phase (OD 0.6) with 1 mM IPTG then incubated overnight at 25 °C to allow protein expression before being harvested by centrifugation at 4,000 g for 15 min at 4 °C. Collected pellets were treated and purified using Ni-NTA affinity chromatography as previously described (26). The T331I mutant was further purified using size exclusion chromatography on a HiLoad 16/600 Superdex 200 (Cytiva) column using a buffer of 50 mM Tris HCl pH 8, 150 mM NaCl, 3 mM TCEP. Fractions containing the target protein were pooled and concentrated using Amicon Ultra Centrifugal Filters. The concentration of the protein was determined by 660 nm protein assay (Pierce) using bovine serum albumin as the reference standard.

### Assessment of autophosphorylation rate and Michaelis-Menten kinetics

The reaction rates of VraS and the T331I mutant were tested using a kinetic coupled assay that measures the rate of ATP consumption per published protocols (26). Briefly, the kinase reaction and conversion of ATP to ADP is coupled to a pyruvate kinase/lactate dehydrogenase (PK/LDH) system that converts NADH to NAD^+^ where the signal monitored was the decrease in NADH absorbance at 340 nm. The reaction rate was monitored over 15 min, and the early linear slope of the reaction (theoretical 10% substrate consumption) was measured. The slopes were mathematically transformed to NADH concentration using the NADH extinction coefficient (6220 L mol^-1^ cm^-1^). The Michaelis Menten kinetic constants for the enzymes were assessed using variable concentrations of ATP as previously described (25).

Construction of T331I *S. aureus* mutant strain.

The C992T point mutation in the *vraS* gene was introduced by recombineering coupled with CRISPR-Cas9 counterselection techniques using published protocols (27, 50). Briefly, a spacer sequence for sgRNA, including the *vraS* 992 position, was selected near a specific protospacer adjacent motif (5’-NGG-3’) site. The annealed 20bps spacer sgRNA with *BsaI* sticky ends (oligos 5&6) was inserted into the pCAS9counter vector, followed by transformation into DH5α. The successful insertion was confirmed by PCR using primers 7&8 (Table S3) followed by Sanger sequencing. To bypass the *S. aureus* restriction barrier, the resulting plasmid pCAS9counter-T331I was further passed through *E. coli* IM08B strain. A ssDNA of 90 bps recombineering oligo 9 with four phosphorothioate bonds at the 5’ end and T331I alteration along with four more wobble positions was designed. The wobble position changes are silent mutations in the protein-coding sequence to avoid bacterial mismatch repair system. The *S. aureus* Newman D2C strain was turned electrocompetent per published protocols and electroporated with the temperature-sensitive recombineering vector pCN-EF2132tet. The resulting strain, Newman pTet, was turned electrocompetent and used to concurrently electroporate the counterselection vector pCAS9counter-T331I and ssDNA of 90 bps oligonucleotide. Positive transformants were selected from LB agar plates containing chloramphenicol and erythromycin (10 μg/mL each). Screening for mutation was conducted by amplifying the *vraS* gene using primers 10&11, and the mutation was confirmed by Sanger sequencing of the amplicon. After sequence confirmation, an isogenic mutant was obtained by streaking on an LB plate (without antibiotics) at 43 °C overnight to cure the temperature-sensitive plasmids. The primer sequences in Table S2, bacterial strains in Table S3, and plasmids in Table S4.

### Drug susceptibility testing

We used the broth microdilution method to determine the MIC of vancomycin, methicillin, and daptomycin (supplemented with 50mg/L CaCl_2_) against the WT and mutant strains using two different growth media CAMHB and LB based on guidelines by the European Committee on Antimicrobial Susceptibility Testing as previously described (25). Briefly, the strains were grown to logarithmic phase in the media, turbidity was adjusted to 0.5 McFarland standard, followed by 1000-fold serial dilution to prepare the inoculum with the bacterial density of ∼10^5^ CFU/ mL. The cultures were co-incubated with the antibiotics at a final drug concentration ranging between 0.25–32 mg/L. At the end of 24hrs of incubation at 37^0^C, plates were visually inspected and the drug concentrations completely inhibiting the bacterial growth was recorded as MIC.

### Concentration- response and growth kinetics studies

We performed dose-response study with both strains in CAMHB, in a total volume of 5 mL. The drug concentrations and the inoculum preparation were the same as described above. After 24hrs of co-incubation with the drugs at 37^0^C under shaking conditions, the cultures were washed twice with normal saline to remove the carry-over drug, serially diluted, and spread on Muller Hinton agar. Cultures were incubated for 24hrs before the CFU/mL with each drug concentration was recorded. GraphPad Prism (V10) was used for data analysis. Growth rates of the WT and T331I mutant strains were determined in LB. Cells were incubated at 37°C with constant shaking in a microplate reader (Synergy H1, BioTek, USA) and the OD600 was measured for 24 hours at 30-minute intervals as previously described (25). All experiments were performed twice in replicates of three.

### Quantitative real-time RT-PCR (qRT-PCR)

The strains were cultured to an OD600 of 0.6 then either LB or selected antibiotics at 2X MIC level were added for 1 hour before harvesting. Total RNA extraction was carried out and the extracted RNA was used for cDNA synthesis to conduct qRT-PCR for target genes per published protocols (25). The data was normalized to the threshold cycle (Ct) value of housekeeping gene 50S ribosomal protein L4 (*rplD*), and the genes’ expression level was obtained using the 2^−ΔΔ*C*t^ method (primer sequences listed in Table S2). Statistical analyses were performed using Prism (V10) using 2-way ANOVA with post-hoc multiple comparisons to determine the statistical significance.

### RNA-seq analysis

The total extracted RNA from four samples (i) New_Van (WT with vancomycin), (ii) T331I_Van (mutant with vancomycin), (iii) New (WT control), and (iv) T331I (control) was sent to Novogene (Sacramento, CA) for DGE analysis. The fragments per kilobase of exon model per million mapped fragments (FPKM) of each gene was calculated using the Feature Counts v1.5.0- p3 for quantitative analysis by mapping to Newman’s genome as a reference. The DGE analysis was performed using Edge R software and statistical tests were performed to identify genes with significant up- or down-regulation compared to the Newman WT strain. The resulting P-values were adjusted using Benjamini and Hochberg’s approach to control the false discovery rate. The p-value < 0.05 and |log2 (Fold change)| value > 1 were set as the threshold for significantly differential expression. For qRT-PCR validation of the top dysregulated genes, the primers were designed using the PrimerQuest™ tool from Qiagen and validated to ensure specificity by ensuring single peaks are obtained in the melt curve and confirmatory PCR and DNA gel resulting in a single product per reaction (Figure S5A). Different primer concentrations (1, 10, 30, and 100 µM) were screened to ensure the sensitivity; the qRT-PCR cycle threshold (Ct) was the lowest in the 10 – 100 µM range. To ensure linear detection, calibration curves were built by screening different concentrations of template cDNA (0.5, 1, 10, and 100 ng/µL) with 10 µM primers. The coefficient of determination (R2) for tested primers was >0.92, indicating good reliability (Figure S5B).

## Acknowledgments

This work was supported by the Fisch College of Pharmacy and UT Tyler Faculty Research Grants, NIH R21AI154189 to M.H.A., and UT Tyler salary support to L.A. The pCAS9counter and pCN-EF2132tet plasmids were a gift from Steve Salipante (Addgene plasmid # 107192 and 107191). The following reagent was obtained through BEI Resources, NIAID, NIH: *Escherichia coli* K-12, Strain IM08B, NR-49806.

## References

1. Turner NA, Sharma-Kuinkel BK, Maskarinec SA, Eichenberger EM, Shah PP, Carugati M, Holland TL, Fowler VG. 2019. Methicillin-resistant Staphylococcus aureus: an overview of basic and clinical research. Nature Reviews Microbiology 2019 17:4 17:203–218.

2. Samia NI, Robicsek A, Heesterbeek H, Peterson LR. 2022. Methicillin-resistant staphylococcus aureus nosocomial infection has a distinct epidemiological position and acts as a marker for overall hospital-acquired infection trends. Scientific Reports 2022 12:1 12:1–10.

3. 2022 SPECIAL REPORT: COVID-19 U.S. Impact on Antimicrobial Resistance | Enhanced Reader. Retrieved 28 October 2024.

4. Liu C, Bayer A, Cosgrove SE, Daum RS, Fridkin SK, Gorwitz RJ, Kaplan SL, Karchmer AW, Levine DP, Murray BE, Rybak MJ, Talan DA, Chambers HF. 2011. Clinical Practice Guidelines by the Infectious Diseases Society of America for the Treatment of Methicillin-Resistant Staphylococcus aureus Infections in Adults and Children. Clinical Infectious Diseases 52:e18–e55.

5. Wielders CLC, Fluit AC, Brisse S, Verhoef J, Schmitz FJ. 2002. mecA Gene Is Widely Disseminated in Staphylococcus aureus Population. J Clin Microbiol 40:3970.

6. Mwangi MM, Shang WW, Zhou Y, Sieradzki K, De Lencastre H, Richardson P, Bruce D, Rubin E, Myers E, Siggia ED, Tomasz A. 2007. Tracking the in vivo evolution of multidrug resistance in Staphylococcus aureus by whole-genome sequencing. Proc Natl Acad Sci U S A 104:9451–9456.

7. Rishishwar L, Petit RA, Kraft CS, Jordana IK. 2014. Genome sequence-based discriminator for vancomycin-intermediate Staphylococcus aureus. J Bacteriol 196:940–948.

8. Cui L, Neoh HM, Shoji M, Hiramatsu K. 2009. Contribution of vraSR and graSR point mutations to vancomycin resistance in vancomycin-intermediate Staphylococcus aureus. Antimicrob Agents Chemother 53:1231–1234.

9. Sharkey LKR, Guerillot R, Walsh CJ, Turner AM, Lee JYH, Neville SL, Klatt S, Baines SL, Pidot SJ, Rossello FJ, Seemann T, McWilliam HEG, Cho E, Carter GP, Howden BP, McDevitt CA, Hachani A, Stinear TP, Monk IR. 2023. The two-component system WalKR provides an essential link between cell wall homeostasis and DNA replication in Staphylococcus aureus. mBio 14.

10. Hafer C, Lin Y, Kornblum J, Lowy FD, Uhlemann AC. 2012. Contribution of Selected Gene Mutations to Resistance in Clinical Isolates of Vancomycin-Intermediate Staphylococcus aureus. Antimicrob Agents Chemother 56:5845.

11. Howden BP, Davies JK, Johnson PDR, Stinear TP, Grayson ML. 2010. Reduced vancomycin susceptibility in Staphylococcus aureus, including vancomycin-intermediate and heterogeneous vancomycin-intermediate strains: Resistance mechanisms, laboratory detection, and clinical implications. Clin Microbiol Rev 23:99–139.

12. Baseri N, Najar-Peerayeh S, Bakhshi B. 2021. Investigating the effect of an identified mutation within a critical site of PAS domain of WalK protein in a vancomycin- intermediate resistant Staphylococcus aureus by computational approaches. BMC Microbiol 21:1–11.

13. Doddangoudar VC, O’donoghue MM, Chong EYC, Tsang DNC, Boost M V. 2012. Role of stop codons in development and loss of vancomycin non-susceptibility in methicillin- resistant Staphylococcus aureus. Journal of Antimicrobial Chemotherapy 67:2101–2106.

14. Gómez-Arrebola C, Hernandez SB, Culp EJ, Wright GD, Solano C, Cava F, Lasa I. 2023. Staphylococcus aureus susceptibility to complestatin and corbomycin depends on the VraSR two-component system. Microbiol Spectr 11:e00370–23.

15. Kuroda M, Kuroda H, Oshima T, Takeuchi F, Mori H, Hiramatsu K. 2003. Two-component system VraSR positively modulates the regulation of cell-wall biosynthesis pathway in Staphylococcus aureus. Mol Microbiol 49:807–821.

16. Gardete S, Wu SW, Gill S, Tomasz A. 2006. Role of VraSR in antibiotic resistance and antibiotic-induced stress response in Staphylococcus aureus. Antimicrob Agents Chemother 50:3424–3434.

17. Dai Y, Gao C, Chen L, Chang W, Yu W, Ma X, Li J. 2019. Heterogeneous vancomycin- intermediate staphylococcus aureus uses the vrasr regulatory system to modulate autophagy for increased intracellular survival in macrophage-like cell line RAW264.7. Front Microbiol 10:452547.

18. Chen H, Xiong Z, Liu K, Li S, Wang R, Wang X, Zhang Y, Wang H. 2016. Transcriptional profiling of the two-component regulatory system VraSR in Staphylococcus aureus with low-level vancomycin resistance. Int J Antimicrob Agents 47:362–367.

19. Su J, Iehara M, Yasukawa J, Matsumoto Y, Hamamoto H, Sekimizu K. 2015. A novel mutation in the vraS gene of Staphylococcus aureus contributes to reduce susceptibility against daptomycin. The Journal of Antibiotics 2015 68:10 68:646–648.

20. Cui L, Neoh HM, Shoji M, Hiramatsu K. 2009. Contribution of vraSR and graSR Point Mutations to Vancomycin Resistance in Vancomycin-Intermediate Staphylococcus aureus. Antimicrob Agents Chemother 53:1231.

21. Katayama Y, Murakami-Kuroda H, Cui L, Hiramatsu K. 2009. Selection of Heterogeneous Vancomycin-Intermediate Staphylococcus aureus by Imipenem. Antimicrob Agents Chemother 53:3190.

22. Sabat AJ, Tinelli M, Grundmann H, Akkerboom V, Monaco M, Grosso M Del, Errico G, Pantosti A, Friedrich AW. 2018. Daptomycin resistant staphylococcus aureus clinical strain with novel non-synonymous mutations in the mprF and vraS genes: A new insight into daptomycin resistance. Front Microbiol 9:411098.

23. Belcheva A, Golemi-Kotra D. 2008. A Close-up View of the VraSR Two-component System: A MEDIATOR OF STAPHYLOCOCCUS AUREUS RESPONSE TO CELL WALL DAMAGE. Journal of Biological Chemistry 283:12354–12364.

24. Galbusera E, Renzoni A, Andrey DO, Monod A, Barras C, Tortora P, Polissi A, Kelley WL. 2011. Site-specific mutation of Staphylococcus aureus VraS reveals a crucial role for the VraR-VraS sensor in the emergence of glycopeptide resistance. Antimicrob Agents Chemother 55:1008–1020.

25. Bhattarai S, Marsh L, Knight K, Ali L, Gomez A, Sunderhaus A, Abdel Aziz MH. 2023. NH125 Sensitizes Staphylococcus aureus to Cell Wall-Targeting Antibiotics through the Inhibition of the VraS Sensor Histidine Kinase. Microbiol Spectr 11.

26. Sunderhaus A, Imran R, Enoh E, Adedeji A, Obafemi T, Abdel Aziz MH. 2022. Comparative expression of soluble, active human kinases in specialized bacterial strains. PLoS One 17:e0267226.

27. Penewit K, Holmes EA, McLean K, Ren M, Waalkes A, Salipante SJ. 2018. Efficient and scalable precision genome editing in Staphylococcus aureus through conditional recombineering and CRISPR/Cas9-mediated counterselection. mBio 9.

28. Drlica K. 2003. The mutant selection window and antimicrobial resistance. Journal of Antimicrobial Chemotherapy 52:11–17.

29. Cutrona N, Gillard K, Ulrich R, Seemann M, Miller HB, Blackledge MS. 2019. From Antihistamine to Anti-infective: Loratadine Inhibition of Regulatory PASTA Kinases in Staphylococci Reduces Biofilm Formation and Potentiates β-Lactam Antibiotics and Vancomycin in Resistant Strains of Staphylococcus aureus. ACS Infect Dis 5:1397–1410.

30. Kanehisa M, Goto S. 2000. KEGG: Kyoto Encyclopedia of Genes and Genomes. Nucleic Acids Res 28:27–30.

31. Gustavsen JA, Pai S, Isserlin R, Demchak B, Pico AR. 2019. RCy3: Network biology using Cytoscape from within R. F1000Res 8:1774.

32. Chen FJ, Lauderdale TL, Lee CH, Hsu YC, Huang IW, Hsu PC, Yang CS. 2018. Effect of a Point Mutation in mprF on Susceptibility to Daptomycin, Vancomycin, and Oxacillin in an MRSA Clinical Strain. Front Microbiol 9:1086.

33. Hiramatsu K, Aritaka N, Hanaki H, Kawasaki S, Hosoda Y, Hori S, Fukuchi Y, Kobayashi I. 1997. Dissemination in Japanese hospitals of strains of Staphylococcus aureus heterogeneously resistant to vancomycin. Lancet 350:1670–1673.

34. Kato Y, Suzuki T, Ida T, Maebashi K. 2010. Genetic changes associated with glycopeptide resistance in Staphylococcus aureus: predominance of amino acid substitutions in YvqF/VraSR. Journal of Antimicrobial Chemotherapy 65:37.

35. Kang YR, Chung DR, Ko JH, Huh K, Cho SY, Kang CI, Peck KR. 2023. Genetic alterations in methicillin-susceptible Staphylococcus aureus associated with high vancomycin minimum inhibitory concentration. Int J Antimicrob Agents 62:106971.

36. Ali L, Aziz MHA. 2024. Crosstalk involving two-component systems in Staphylococcus aureus signaling networks. J Bacteriol 206.

37. Trajtenberg F, Graña M, Ruétalo N, Botti H, Buschiazzo A. 2010. Structural and Enzymatic Insights into the ATP Binding and Autophosphorylation Mechanism of a Sensor Histidine Kinase. Journal of Biological Chemistry 285:24892–24903.

38. Landry BP, Palanki R, Dyulgyarov N, Hartsough LA, Tabor JJ. 2018. Phosphatase activity tunes two-component system sensor detection threshold. Nature Communications 2018 9:1 9:1–10.

39. Brouwers R, Vass H, Dawson A, Squires T, Tavaddod S, Allen RJ. 2020. Stability of β- lactam antibiotics in bacterial growth media. PLoS One 15:e0236198.

40. Lade H, Kim JS. 2023. Molecular Determinants of β-Lactam Resistance in Methicillin-Resistant Staphylococcus aureus (MRSA): An Updated Review. Antibiotics 2023, Vol 12, Page 1362 12:1362.

41. Muthaiyan A, Silverman JA, Jayaswal RK, Wilkinson BJ. 2008. Transcriptional profiling reveals that daptomycin induces the Staphylococcus aureus cell wall stress stimulon and genes responsive to membrane depolarization. Antimicrob Agents Chemother 52:980–990.

42. Hort M, Bertsche U, Nozinovic S, Dietrich A, Schrötter AS, Mildenberger L, Axtmann K, Berscheid A, Bierbaum G. 2021. The Role of β-Glycosylated Wall Teichoic Acids in the Reduction of Vancomycin Susceptibility in Vancomycin-Intermediate Staphylococcus aureus. Microbiol Spectr 9.

43. Berscheid A, François P, Strittmatter A, Gottschalk G, Schrenzel J, Sass P, Bierbaum G. 2014. Generation of a vancomycin-intermediate Staphylococcus aureus (VISA) strain by two amino acid exchanges in VraS. Journal of Antimicrobial Chemotherapy 69:3190– 3198.

44. Lee JJ, Lee SK, Song N, Nathan TO, Swarts BM, Eum SY, Ehrt S, Cho SN, Eoh H. 2019. Transient drug-tolerance and permanent drug-resistance rely on the trehalose-catalytic shift in Mycobacterium tuberculosis. Nature Communications 2019 10:1 10:1–12.

45. Dubrac S, Boneca IG, Poupel O, Msadek T. 2007. New insights into the WalK/WalR (YycG/YycF) essential signal transduction pathway reveal a major role in controlling cell wall metabolism and biofilm formation in Staphylococcus aureus. J Bacteriol 189:8257– 8269.

46. Lopes AA, Yoshii Y, Yamada S, Nagakura M, Kinjo Y, Mizunoe Y, Okuda Kichi. 2019. Roles of Lytic Transglycosylases in Biofilm Formation and β-Lactam Resistance in Methicillin- Resistant Staphylococcus aureus. Antimicrob Agents Chemother 63.

47. Audretsch C, Gratani F, Wolz C, Dandekar T. 2021. Modeling of stringent-response reflects nutrient stress induced growth impairment and essential amino acids in different Staphylococcus aureus mutants. Scientific Reports 2021 11:1 11:1–15.

48. Boyle-Vavra S, Yin S, Jo DS, Montgomery CP, Daum RS. 2013. VraT/YvqF is required for methicillin resistance and activation of the VraSR regulon in Staphylococcus aureus. Antimicrob Agents Chemother 57:83–95.

49. Łȩski TA, Tomasz A. 2005. Role of penicillin-binding protein 2 (PBP2) in the antibiotic susceptibility and cell wall cross-linking of Staphylococcus aureus: Evidence for the cooperative functioning of PBP2, PBP4, and PBP2A. J Bacteriol 187:1815–1824.

50. Penewit K, Salipante SJ. 2020. Genome Editing in Staphylococcus aureus by Conditional Recombineering and CRISPR/Cas9-Mediated Counterselection. Methods in Molecular Biology 2050:127–143.

